# Organoid-derived Duodenum Intestine-Chip for preclinical drug assessment in a human relevant system

**DOI:** 10.1101/723015

**Authors:** Magdalena Kasendra, Raymond Luc, Jianyi Yin, Dimitris V. Manatakis, Athanasia Apostolou, Laxmi Sunuwar, Jenifer Obrigewitch, Geraldine A. Hamilton, Mark Donowitz, Katia Karalis

**Affiliations:** Emulate Inc., 27 Drydock Avenue, Boston, MA 02210, USA; Department of Medicine, Division of Gastroenterology, Johns Hopkins University School of Medicine, Baltimore, MD 21205, USA; Graduate Program, Department of Medicine, National and Kapodistrian University of Athens, Greece

## Abstract

Induction of intestinal drug metabolizing enzymes can complicate the development of new drugs, owing to potential to cause drug-drug interactions (DDIs) leading to changes in pharmacokinetics, safety and efficacy. The development of a human relevant model of the adult intestine that accurately predicts CYP450 induction could help address this challenge as species differences preclude extrapolation from animals. Here, we combined organoids and Organ-Chip technology to create a human Duodenum Intestine-Chip that emulates intestinal tissue architecture and functions, that are relevant for the study of drug transport, metabolism, and DDI. Duodenum Intestine-Chip demonstrates the polarized cell architecture, intestinal barrier function, presence of specialized cell subpopulations, and *in vivo*-relevant expression, localization, and function of major intestinal drug transporters. Notably, in comparison to Caco-2, it displays improved CYP3A4 expression and induction capability. This model could enable improved *in vitro* to *in vivo* extrapolation for better predictions of human pharmacokinetics and risk of DDIs.

## Introduction

A challenge in drug development is low oral bioavailability and poor pharmacokinetics caused by drug-drug interactions of orally administered drug. One key driver can be high affinity for drug transporters and activity of metabolic enzymes in the human intestine ^1–4^. After the decades of research, the hepatic drug clearance is well-understood and relatively well predicted by pre-clinical models. While accurate prediction of first-pass extraction of xenobiotics in human intestinal epithelium still remains elusive. This is due to a number of confounding factors that affect oral drug absorption including the properties of the compound (solubility, permeability), physiology of the intestinal tract (transit time, blood flow), and patient phenotype (including age, gender, polymorphism in drug metabolizing enzymes, disease states) ^5^. Species differences in the isoforms, regional abundances, differences in substrate specificity of drug metabolism enzymes ^6–8^ and transporters ^9, 10^, and mechanism regulating transcriptional activation ^11, 12^, precludes accurate extrapolation of the data from animal models to the clinic. In mice, for example, there are 34 cytochromes CY450 in the major gene families involved in drug metabolism, i.e., the Cyp1a, Cyp2c, Cyp2d, and Cyp3a gene clusters, while in humans, there are only eight^13^. Interestingly three human enzymes, CYP2C9, CYP2D6, and CYP3A4, account for ∼75% of all reactions, with CYP3A4 being the single most important human CYP450 accounting for ∼45% of phase 1 drug metabolism in humans ^14^. In addition, the expression levels of many of the major human CYP450 enzymes and drug transporter (which determine levels and variability in drug exposure) are controlled by multiple transcription factors, primarily the xenosensors constitutive androstane receptor (CAR), pregnane X receptor (PXR), and aryl hydrocarbon receptor (AhR). These nuclear receptors also exhibit marked species differences in their activation by drugs and exogenous chemicals ^11^. For example, rifampicin and SR12813 are potent agonists for human PXR (hPXR) but not for mouse PXR (mPXR), whereas the potent mPXR agonist 5-pregnen-3β-ol-20-one-16α-carbonitrile (PCN) is a poor agonist for hPXR ^15^. On the other hand, 6-(4-chlorophenyl)imidazo[2,1-b][1,3]thiazole-5-carbaldehyde-O-(3,4-dichlorobenzyl)oxime (CITCO) is a strong agonist for human CAR (hCAR) but not mouse CAR (mCAR) ^16^, while 1,4-bis-[2-(3,5-dichloropyridyloxy)]benzene,3,3′,5,5′-tetrachloro-1,4-bis(pyridyloxy)benzene (TCPOBOP) is more selective for mCAR than hCAR. Such species differences together with the complex interplay between drug metabolizing enzymes and drug transporters in the intestine and the liver, as well as, the overlap of substrate and inhibitor specificity ^17^, make it difficult to predict human pharmacokinetics when there is intestinal metabolism and drug transport involved due to the lack of predictive pre-clinical models that reflect the human biology that allow researchers to study it in a human relevant system.

Numerous *in vitro* systems have been developed and applied routinely for characterization of distribution and prediction of absorption, distribution, metabolism, and excretion (ADME) of potential drug candidates in humans. Among these is Caco-2 monolayer culture on a transwell insert, which is the one of most widely used across the pharmaceutical industry as an *in vitro* representation of the human small intestine. However, inherent limitations, such as lack of *in vivo* relevant three-dimensional cytoarchitecture, lack of appropriate ratio of cell populations, altered expression profiles of drug transporters and drug metabolizing enzymes, especially CYP450s, and aberrant CYP450 induction response, challenge the use of these model for predicting human responses in the clinic ^18^.

A promising alternative to conventional cell monolayer systems emerged with the establishment of the protocols for generation of three-dimensional intestinal organoids (or enteroids) from human biopsy specimens ^19–22^. Using these methods, organoids derived from all regions of the intestinal tract can be established ^21, 23^ and applied into different areas of research including organ development, disease modeling, and regenerative medicine ^24^. However, the characterization of the pharmacokinetic properties of this system, as well as the its validation for the use in drug discovery and development, is still very limited ^25–29^. One potential limitation could be due to the substantial technical challenges associated with the use of this technology for ADME applications. The 3D-spheroidal architecture of the organoid restricts access to its lumen, which is crucial for assessing intestinal permeability or drug absorption. Indeed, exposure of the apical cell surface of intestinal organoid to compounds requires the use of time-consuming and labor-intensive procedures, such as microinjection. The presence of relatively thick gel of extracellular matrix (Matrigel^TM^) surrounding organoids, might limit drug penetration. While heterogeneity of organoids in terms of their size, shape, and viability, can also impede studies in ADME and robust results ^24^.

In addition, none of these models have fully recapitulated critical aspects of an organ microenvironment such as the presence of microvasculature, mechanical forces of fluid flow (shear stress) and peristalsis, all of which contribute to capturing the complex and dynamic nature of *in vivo* tissue function ^30^. For these reasons, there is a need for new systems for predicting human ADME and determining risk for drug-drug interactions mediated by intestinal CYP450s and drug transporters in the clinic.

We have recently developed a human Duodenum Intestine-Chip that combines healthy intestinal organoids with our Organs-on-Chips technology ^31^ to overcome the existing limitations of the current systems. Here, we demonstrated that the Duodenum Intestine-Chip provides a more human-relevant model, in respect to organoids, as supported by the comparison of their global RNA gene expression profiles, and that it can be applied for the study of drug transport, drug metabolism, and drug-drug interactions. Presence of the mechanical forces, which are applied into this system in order to recapitulate the blood flow and shear stress, showed to improve the formation of the right cell cytoarchitecture and the increased appearance of intestinal microvilli on the apex cell surface. In addition, Duodenum Intestine-Chip supported successful maturation of all major intestinal cell types in the physiologically relevant ratios and low paracellular permeability. Importantly for its application into pharmacokinetic studies, it showed closer to *in vivo* expression of drug uptake and efflux transporters, as compared to Caco-2-cells based system. Additionally, we were able to prove that it the system supports the correct luminal localization and functional activity of MDR1 (P-gp), as well as high expression of cytochrome CYP450 (CYP) 3A4 - close to the levels observed in human duodenal tissue. Importantly, exposure of the Duodenum Intestine-Chip to known CYP450 inducers in humans, such as rifampicin and 1, 25-dihydroxyvitamin D3, resulted in the significant increase of both mRNA and protein levels of CYP3A4. Our results indicate that the organoid-derived human tissue combined with Organs-on-Chips technology could provide a robust human-relevant system the assessment of CYP450 metabolism, drug transporter effects, and risk for potential drug-drug interactions in preclinical testing.

### Development of the adult Duodenum Intestine-Chip

We have previously developed a human organoid-based Intestine-Chip, referred to at the time as the “Small Intestine-on-a-Chip”, which combined the use of intestinal organoids and Organ-Chips ^32^ and provides a system to model intestinal biology. Here we sought to establish the adult Duodenum Intestine-Chip to serve as a model for preclinical assessment of drug transport and metabolism. In brief, we first established cultures of organoids (Figure 1A; top) derived from the endoscopic biopsies of three different healthy adult donors; the organoids were then dissociated into fragments and seeded on the ECM-coated porous flexible poly(dimethylsiloxane) (PDMS) membrane of the chips (Figure 1B; 1: indicates the epithelial tissue). Primary human intestinal microvascular endothelial cells (HIMECs, Cell Biologics), derived from the human small intestine (Figure 1A; bottom), were used to populate the other surface of the PDMS membrane in the vascular channel (Figure 1B; 5: indicates the endothelial cells). Next, the Duodenum Intestine-Chips were perfused continuously through luminal and vascular compartment with 30 µl / hr of fresh media. Once the epithelial monolayers reached confluency they were subjected to cyclic mechanical strain (10% strain, 0.2 Hz) in order to emulate intestinal peristalsis and the physiologically relevant mechanical forces associated with it.

**Figure 1.**
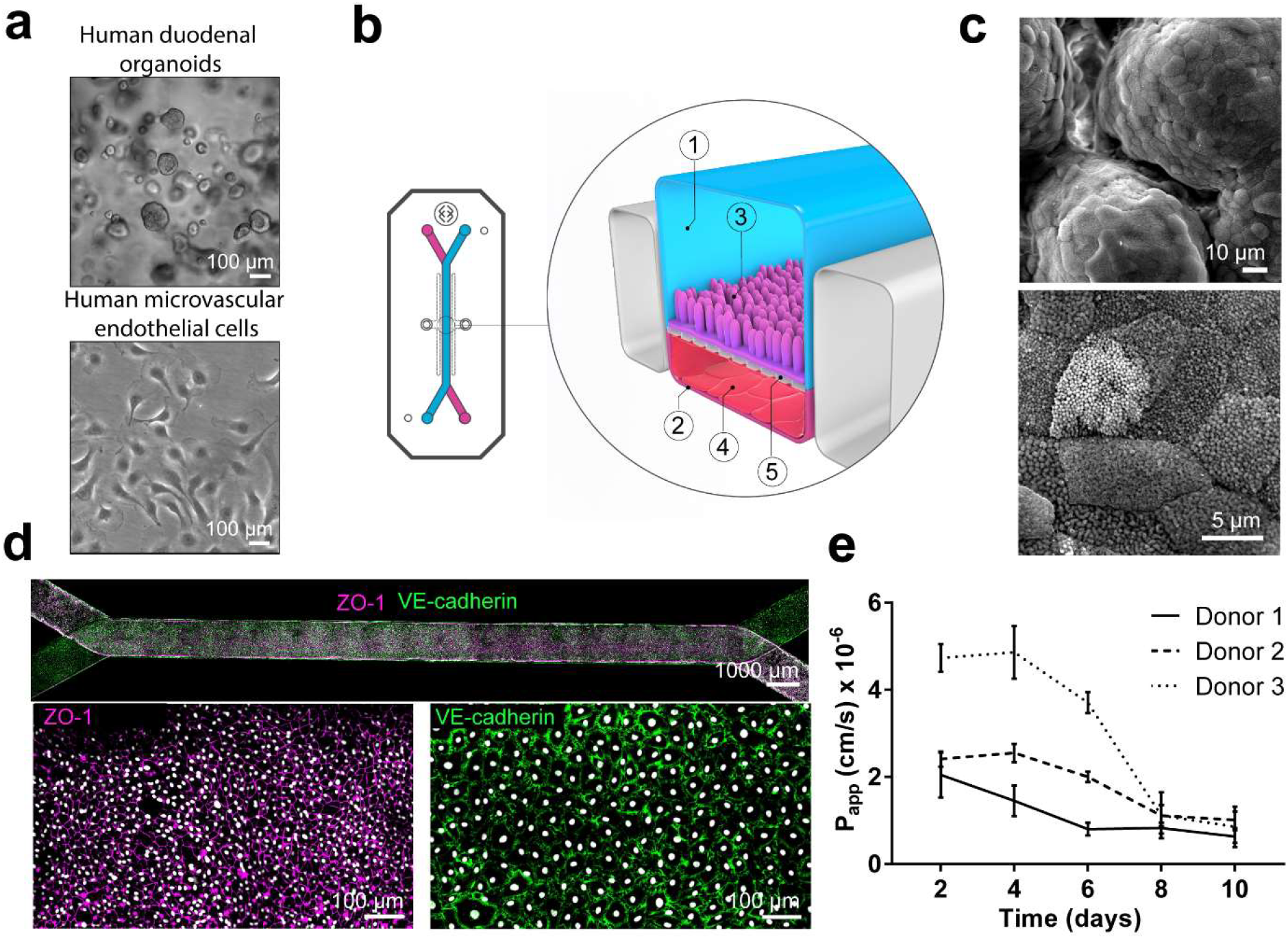
Duodenum Intestine-Chip: a microengineered model of the human duodenum. **(a)** Brightfield images of human duodenal organoids (top) and human microvascular endothelial cells (bottom) acquired before their seeding into epithelial and endothelial channels of the chip. **(b)** Schematic representation of Duodenum Intestine-Chip, including its top view (left) and vertical section (right) showing: the epithelial (1; blue) and vascular (2; pink) cell culture microchannels populated by intestinal epithelial cells (3) and endothelial cells (4), respectively, and separated by a flexible, porous, ECM-coated PDMS membrane (5). **(c)** Scanning electron micrograph showing complex intestinal epithelial tissue architecture achieved by duodenal epithelium grown on the chip (top) in the presence of constant flow of media (30 µl h^−1^) and cyclic membrane deformations (10% strain, 0.2 Hz). High magnification of the apical epithelial cell surface with densely packed intestinal microvilli (bottom). **(d)** Composite tile scan fluorescence image (top) showing a fully confluent monolayer of organoids-derived intestinal epithelial cells (magenta, ZO-1 staining) lining the lumen of Duodenum Intestine-Chip and interfacing with microvascular endothelium (green, VE-cadherin staining) seeded in the adjacent vascular channel. Higher magnification views of epithelial tight junctions (bottom left) stained against ZO-1 (magenta) and endothelial adherence junctions visualized by VE-cadherin (green) staining. Cells nuclei are shown in grey. Scale bar, 1000 µm (top) 100 µm (bottom) **(e)** Apparent permeability values of Duodenum Intestine-Chips cultured in the presence of flow and stretch (30 ul/h, 10% 0,2 Hz) for up to 10 days. Papp values were calculated from the diffusion of 3 kDa Dextran from the luminal to the vascular channel. Data represent three independent experiments performed with three different chips/donor, total of 3 donors; Error bars indicate s.e.m.

We then assessed the effect of applying mechanical forces (via fluid flow and stretch) on the phenotypic characteristics of the human primary intestinal cells in the chips. We used multiple endpoints including immunofluorescent staining for apical (villin) and basolateral (E-cadherin) cell surface markers and scanning electron microscopy (SEM) for the presence of apical microvilli. In line with our previous findings in the Caco-2 Intestine-Chip^33^, exposure of the Duodenum Intestine-Chip to flow for 72 hours resulted in accelerated cell polarization and formation of apical microvilli (Supplementary Figure 1). We found that cells cultured under static conditions in the Organ-Chips formed a flat (14.8 ± 2.6 μm) monolayer of epithelial cells of squamous appearance and were characterized by poorly defined cell-cell junctions (Supplementary Figure 1A) and sparsely distributed microvilli (Supplementary Figure 1B). In contrast, cells cultured under flow (30 μLh^−1^) with or without concomitant application of cyclic stretch (10%, 0.2 Hz), exhibited a well-polarized and cobblestone-like morphology with increased cell height (27.0 ± 1.3 μm), strongly delineated junctions, and dense microvilli-like structures. In line with our previous findings with Caco-2 cells, the application of constant flow (shear stress) was critical for promoting maturation of a well-polarized epithelium, while short-term application of cyclic strain did not show any additional effect. In addition, more prolonged exposure of cells to flow and cyclic strain resulted in the spontaneous development of epithelial undulations (“villi-like structures”) extending into the lumen of the epithelial channel and covered by continuous brush border (Figure 1C). Immunofluorescence confocal analysis confirmed the establishment of confluent epithelial and endothelial monolayers across the entire length of the chip (Figure 1D), with well-defined epithelial tight junctions, as demonstrated by *in vivo*-relevant ZO-1 protein staining and endothelial adherent junctions visualized using antibody against VE-cadherin (green). Importantly, these culture conditions resulted in a time-dependent improvement of intestinal permeability as indicated by the low permeability coefficient (Papp) of fluorescently labeled dextran recorded in the Duodenum Intestine-Chips generated from 3 different human donors (Figure 1E). This data indicates that this model supports the formation of a functional barrier with *in vivo* relevant cytoarchitecture, cell-cell interactions, and permeability parameters.

To confirm differentiation of the organoid-derived primary enterocytes to the relevant subpopulations of cells as found *in vivo*, we assessed expression levels of cell type specific markers, including alkaline phosphatase (ALPI) for absorptive enterocytes, mucin 2 (MUC2) for goblet cells, chromogranin A (CHGA) for enteroendocrine cells, and lysozyme (LYZ) for Paneth cells in Duodenum Intestine-Chips from 3 different human donors. We compared expression levels of these genes to those found in freshly isolated adult duodenal tissue (Duodenum). As shown (Figure 2A), expression of all the above genes, with the exception of LYZ, was increased over time in culture. Notably, markers such as alkaline phosphatase and mucin 2 reached similar levels in Duodenum Intestine-Chips to those detected in RNA isolated directly from the human duodenum after 8 days in culture. The opposite trend depicted in the expression of LYZ suggests a decrease in Paneth cells in line with increased differentiation of the villus positive epithelium and a decrease in progenitor/stem cells population. In addition, immunostaining for the characterization of all major differentiated intestinal cell types, followed by quantification using confocal microscopy, showed physiologically relevant relative ratios of each cell-type to one another (Figure 2C). The ratios of the cell populations in the chips were close to those reported following histopathological analysis of sections from the human duodenum (Figure 2D)^34^. Taken together, these results demonstrate the successful establishment of an adult Duodenum Intestine-Chip that closely recreates the barrier function and multilineage differentiation of adult human intestinal tissue.

**Figure 2.**
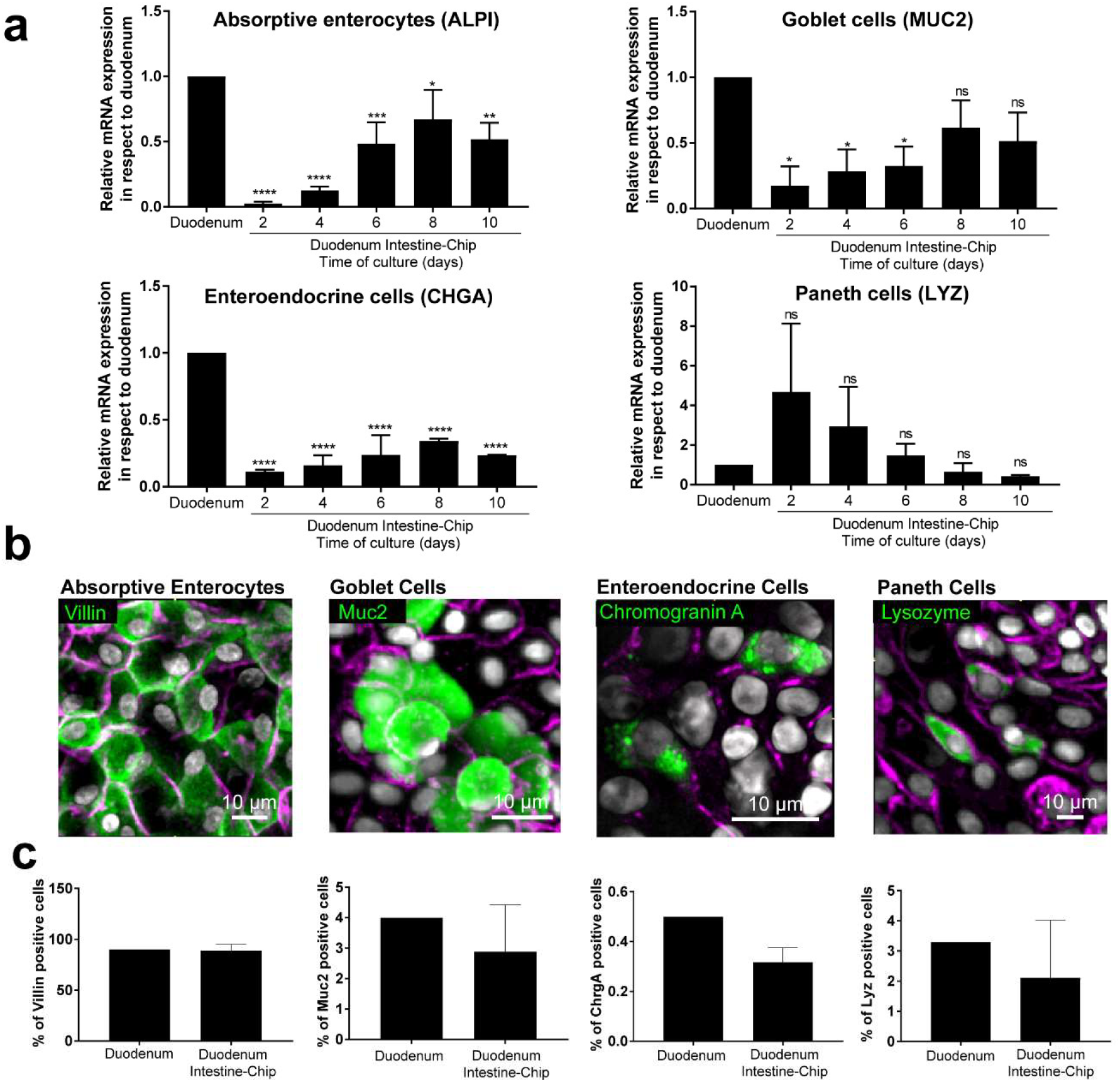
Duodenum Intestine-Chip emulates multi-lineage differentiation of native human intestine. **(a)** Comparison of the relative gene expression levels of markers specific for different intestinal cell types, including mucin 2 (MUC2) for goblet cells, alkaline phosphatase (ALPI) for absorptive enterocytes, chromogranin A (CHGA) for enteroendocrine cells, lysozyme (LYZ) for Paneth cells, leucine-rich repeat-containing G-protein coupled receptor 5 (LGR5) for stem cells and proliferation marker (KI67) across Duodenum Intestine-Chips established from 3 individual organoids donors at different times in the chip under continuous flow (day 2, 4, 6, 8, 10) and RNA isolated directly from the duodenal tissue of other 3 different donors (Duodenum). In each graph, values represent average gene expression ± s.e.m (error bars) from three independent experiments, each using different donors of biopsy-derived organoids and at least three different chips per time point. Values are shown relative to duodenal tissue expressed as 1. EPCAM expression was used as normalizing control. One-way ANOVA, ****p<0.0001, ***p<0.001, **p<0.01, *p<0.05, ns p>0.05 **(b)** Representative confocal fluorescent micrographs demonstrating the presence of all major intestinal cell types (green) in Duodenum Intestine-Chip at day 8 of fluidic culture, including goblet cells stained with anti-Muc2; enteroendocrine cells visualized with anti-chromogranin A, absorptive enterocytes stained with anti-villin and Paneth cells labeled with anti-lysozyme. Cell-cell borders are stained with anti-E-cadherin and are shown in magenta. Scale bar, 10 µm **(c)** Quantification of the different epithelial intestinal cell types present in Duodenum Intestine-Chip at day 8 and identified by immunostaining, as described in (b). Percentage of different cell types based on 10 different fields of view (10 FOV) counted in three individual chips per staining established from three different donors. DAPI staining was used to evaluate the total cell number. Duodenum values, represent cell ratios observed in the histological sections and are based on the literature ^34^.

### Transcriptomic comparison of the Duodenum Intestine-Chip versus organoids

To further verify whether Duodenum Intestine-Chip faithfully recapitulates human adult duodenum tissue and to better understand how much it differs from the organoids from which it’s derived, we performed RNA-seq analysis. We compared global RNA expression data obtained from: i) duodenum organoids (Organoids; n=3) cultured in conventional plastic-adherent Matrigel^TM^ drop overlaid with growth medium; ii) Duodenum Intestine-Chips derived from organoids (Duodenum Intestine-Chip; n=3) in the presence of constant flow and stretch; iii) human adult duodenum tissue (adult duodenum; n=2; full-thickness samples) (Supplementary Table 1). We annotated 13,735 genes in the genome and performed differential gene expression analysis (DGE). For the DGE analysis, we used the “limma” R package and applied the widely accepted thresholds: adjusted p-value<0.05 and |log2FoldChange| > 2 in order to select the differentially expressed genes^35^.

First, we examined the differential gene expression in organoids compared to human adult duodenum tissue. Out of the 13,735 genes annotated in the genome, 1437 were found to be significantly differentially regulated among these samples: 562 and 875 genes were respectively up- and downregulated (Supplementary Figure 2A and Supplementary Table 2). Next a functional enrichment analysis was performed utilizing the PANTHER classification system to highlight biological processes, i.e., significantly enriched gene ontology (GO) terms within these gene sets ^36–38^. The majority of differentially expressed genes belonged to pathways related to digestion, extracellular matrix organization, angiogenesis, cell adhesion, tissue development, and cell responses to drugs and xenobiotics (Supplementary Figure 2A). This comparison allowed us to identify genes responsible for global transcriptomic differences between organoids technology and native human tissue and highlight biological functions which could be affected by the observed differences.

We applied similar type of analysis, to the one described above, to compare global gene expression profile of Duodenum Intestine-Chip with adult duodenum. This analysis resulted in the identification of 1023 differentially expressed genes that were significantly up-(382 genes) or downregulated (641 genes). The fact, that the significantly smaller number of differentially expressed genes was identified in this analysis as compared to the DGE performed between organoids and adult duodenum tissue, suggests that combining organoids with Organs-on-Chips enables a better emulation of the human duodenum tissue (Supplementary Figure 2B and Supplementary Table 3). Functional enrichment analysis highlighted biological processes such as protein synthesis and targeting, cell cycle and cell proliferation underlying the differences between Duodenum Intestine-Chip model and native human tissue.

Next, in order to determine which genes are responsible for a closer similarity of Duodenum Intestine-Chip system to human duodenum tissue in comparison to organoids, we carried out additional DGE analysis. This time we assessed the differences between Duodenum Intestine-Chip and organoids, and examined these differences relative to those that exist when comparing the adult human duodenal tissue and organoids (Duodenum Intestine-Chip versus organoids and human adult duodenum versus organoids). We identified genes that were significantly up- and downregulated in the Duodenum Intestine-Chip relative to organoid culture (Figure 3—source data 1) and identified the proportion of these genes that were also significantly different in adult duodenal tissue relative to organoids (Figure 3—source data 2 and 6). We found 305 genes which were common for Duodenum Intestine-Chip and adult duodenum but different from organoids, representing 39.25% of differentially expressed genes in Duodenum Intestine-Chip versus organoids comparison, and 21.22% of differentially expressed genes in human adult duodenal tissue versus organoids comparison (Figure 3A and Figure 3—source data 3). To further explore overlapping genes, we used the Gene Ontology and Kyoto Encyclopedia of Genes and Genomes (KEGG) Pathway Analysis to examine biological processes and pathways that are enriched in this gene list ^39, 40^. In addition, we used the well-established Reduce and Visualize Gene Ontology (REVIGO) tool to reduce redundancies by clustering semantically related GO terms and outputting them as a scatter plot for visualization ^41, 42^. As a result, the list of 305 overlapping genes was associated with 117 significant GO terms, which were reduced to 74 unique GO terms in REVIGO (Figure 3—source data 4). Notably, biological processes enriched in this gene set were associated with important intestinal functions such as digestion and transport of nutrients and ions, metabolism, detoxification, as well as tissue and tube development (Fig 3B).

**Figure 3.**
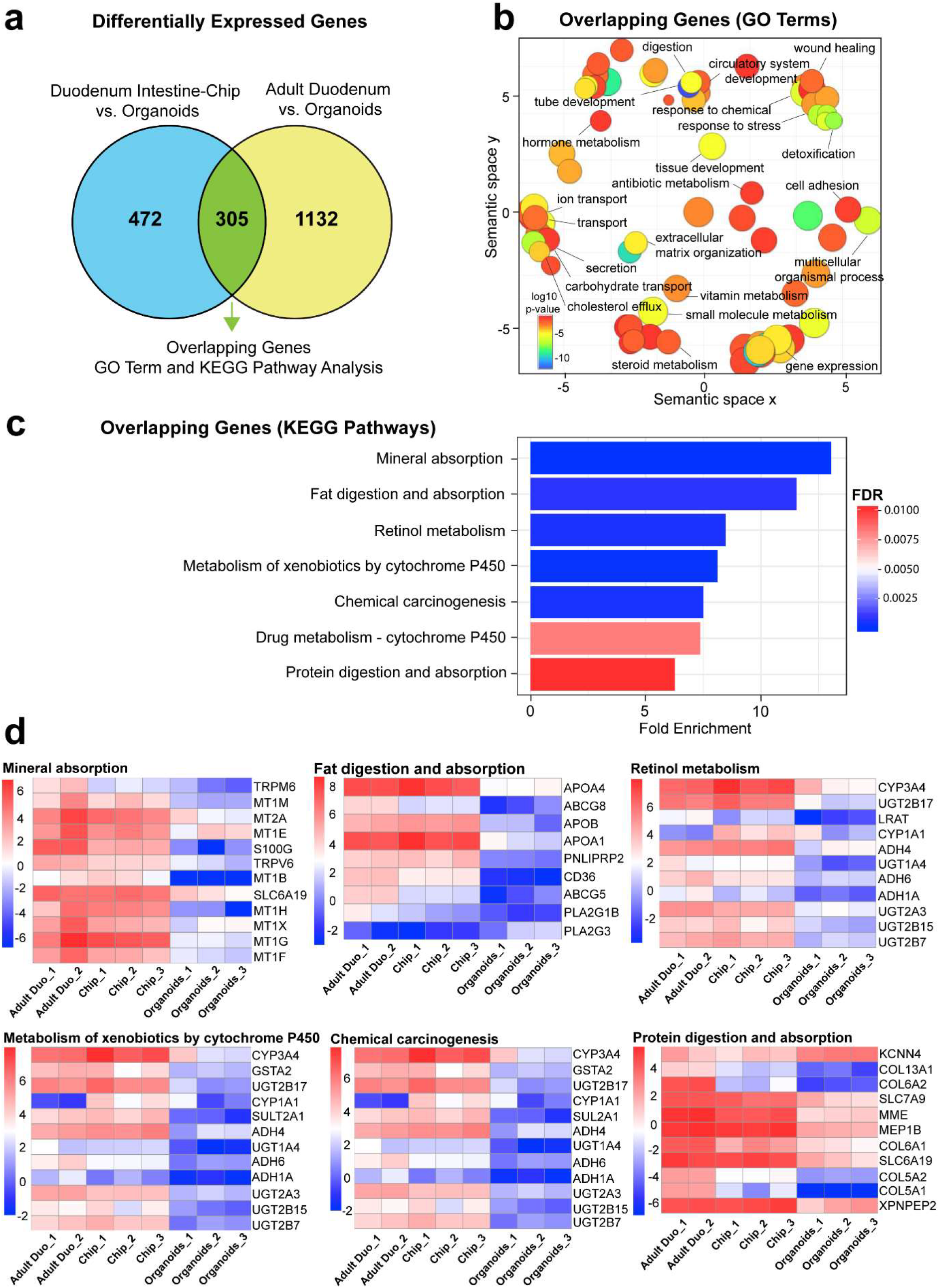
Duodenum Intestine-Chip exhibits higher transcriptomic similarity to adult duodenal tissue than organoid culture. **(a)** Differential gene expression analysis was carried out to identify genes that are upregulated or downregulated in Duodenum Intestine-Chip compared to organoids (blue circle) (Figure 3—source data 1) and adult duodenum compared to organoids (yellow circle) (Figure 3—source data 2). The gene lists were then compared to determine how many genes overlap between those two comparisons (green) (Figure 3—source data 3), and the results are shown as a Venn diagram. 305 genes were identified as common and responsible for the closer resemblance of Duodenum Intestine-Chip to human adult duodenum than organoids from which chips were derived. Sample sizes were as follows: Duodenum Intestine-Chip, n =3 (independent donors); Organoids, n = 3 (independent donors); Adult duodenum, n = 2 (independent biological specimens). Intestinal crypts derived from the same three independent donors were used for the establishment of Duodenum Intestine-Chip and organoid cultures **(b)** The list of overlapping genes was subjected to GO analysis to identify enriched biological processes (GO terms) (Figure 3—source data 4). The results are shown as REVIGO scatterplots in which similar GO terms are grouped in arbitrary two-dimensional space based on semantic similarity. Each circle corresponds to a specific GO term and circle sizes are proportional to the number of genes included in each of the enriched GO terms. Finally, the color of a circle indicates the significance of the specific GO term enrichment. GO terms enriched in the overlapping gene set demonstrate that Duodenum Intestine-Chip is more similar to human duodenum with respect to important biological functions of the intestine, including digestion, transport and metabolism. **(c)** The results of the KEGG pathway analysis using the 305 differentially expressed genes showed seven significantly enriched (FDR corrected p-value <0.05) pathways related to absorption, metabolism, digestion and chemical carcinogenesis. The size of the bars indicates the fold-enrichment of the corresponding pathways. **(d)** Curated heatmaps were generated to examine particular genes that belong to the enriched KEGG pathways and show the expression intensity (log_2_(FPKM) of these genes across different samples. The provided results further demonstrate that Duodenum Intestine-Chip is more similar to adult duodenum than are the organoids. Sample sizes were as follows: Duodenum Intestine-Chip, n =3 (independent donors); Organoids, n = 3 (independent donors); Adult duodenum, n = 2 (independent biological specimens). Intestinal crypts derived from the same three independent donors were used for the establishment of Duodenum Intestine-Chip and organoid cultures. **Figure 3—source data 1.** Differentially Expressed Genes in Duodenum Intestine-Chip vs. Duodenal Organoids, Related to Figure 3. (added as Supplementary Materials) **Figure 3—source data 2.** Differentially Expressed Genes in Adult Duodenum vs. Duodenal Organoids, Related to Figure 3. (added as Supplementary Materials) **Figure 3—source data 3.** Differentially Expressed Genes Common in Duodenum Intestine-Chip and Adult Duodenum versus Duodenal Organoids, Related to Figure 3. (added as Supplementary Materials) **Figure 3—source data 4.** Enriched GO Terms from a List of Differentially Expressed Genes Common in Duodenum Intestine-Chip and Adult Intestine versus Duodenal Organoids, Related to Figure 3. (added as Supplementary Materials)

Additionally, KEGG Pathway analysis identified the total of 7 significantly enriched pathways (threshold FDR p-value <0.05) among the overlapping genes (Figure 3C). The corresponding fold enrichments of these pathways are shown in Figure 3C. Importantly, many of these pathways were linked to drug metabolism, such as – cytochrome CYP450 (associated with 10 genes), metabolism of xenobiotic by cytochrome CYP450 (associated with 12 genes), and other important pathways related to nutrients absorption and digestion, such as mineral absorption (associated with 12 genes), fat digestion and absorption (associated with 9 genes), protein digestion and absorption (associated with 11 genes) (Figure 3C). To more closely examine gene signatures that belong to these pathways, we used log (FPKM) values generated from the RNA-seq dataset, and generated heatmaps for differentially expressed genes that fall under the 7 significantly enriched pathways (Figure 3D). A large number of genes involved in the nutrient absorption, digestion, as well as, metabolism of xenobiotics was highly expressed in Duodenum Intestine-Chip (Chip) and adult duodenum tissue (Adult Duo) but expressed at the low level in organoids.

Cumulatively, these data demonstrated an increased similarity in the global gene expression profile between Duodenum Intestine-Chip and human adult duodenum tissue in comparison to the organoids from the same donor.

### Intestinal drug transporters and MDR1 efflux activity in the Duodenum Intestine-Chip

Encouraged by the transcriptomic analysis detailed above, that showed a clear advantage of Duodenum Intestine-Chip over organoids from which it was derived and a closer to *in vivo* expression of genes related to absorption, transport, and metabolism we next sought to focus further on the expression, localization and function of main intestinal drug transporters. We determined their expression and localization in the Duodenum Intestine-Chips established from the organoids of three independent donors by qRT-PCR and immunofluorescent imaging. We have compared the observed gene expression levels with the values obtained for the freshly isolated human duodenal tissue samples (Duodenum) and the previously described Intestine-Chip model based on the use of Caco-2 cells (Caco-2 Intestine-Chip) (Figure 4A). Averaged gene expression levels of efflux (MDR1, BCRP, MRP2, MRP3) and uptake (PepT1, OATP2B1, OCT1, SLC40A1) drug transporters in the Duodenum Intestine-Chips were close to those observed in the human duodenal tissue. Similar pattern of gene expression was revealed for Caco-2 Intestine-Chip, suggesting the existence of a good correlation between the chip and human *in vivo* tissue. However, in case of the latter model, Caco-2 Intestine-Chip, a couple of important organic anion and cation transporters, including OATP2B1 and OCT1, showed a markedly increased expression in comparison to human duodenum. This confirmed the maintenance of already well-known differences between the cancer-derived cell line and normal human intestinal tissue in Intestine-Chip system^43, 44^. Moreover, although not statistically significant, but notable differences in the expression of MDR1, BCRP and PEPT1 were present in Caco-2 Intestine-Chip in comparison to Duodenum. In line with the trends previously reported by others^43, 45, 46^, a 3.5-fold higher expression of MDR1, 3.6-fold and 24-fold lower expression of BCRP and PEPT1, respectively, were found in Caco-2 cells compared with human duodenum. On the other hand, much smaller differences in respect to native human tissue were observed for Duodenum Intestine-Chip: with ~1.5-fold and ~2.6-fold increased expression of MDR1 and BCRP, respectively, and no differences noted in the expression of PEPT1. These results demonstrate that by combining duodenal organoids with Organ-on-Chips technology we enabled a closer emulation of human *in vivo* tissue than it was possible with the previously described Caco-2 Intestine-Chip or organoids alone.

**Figure 4.**
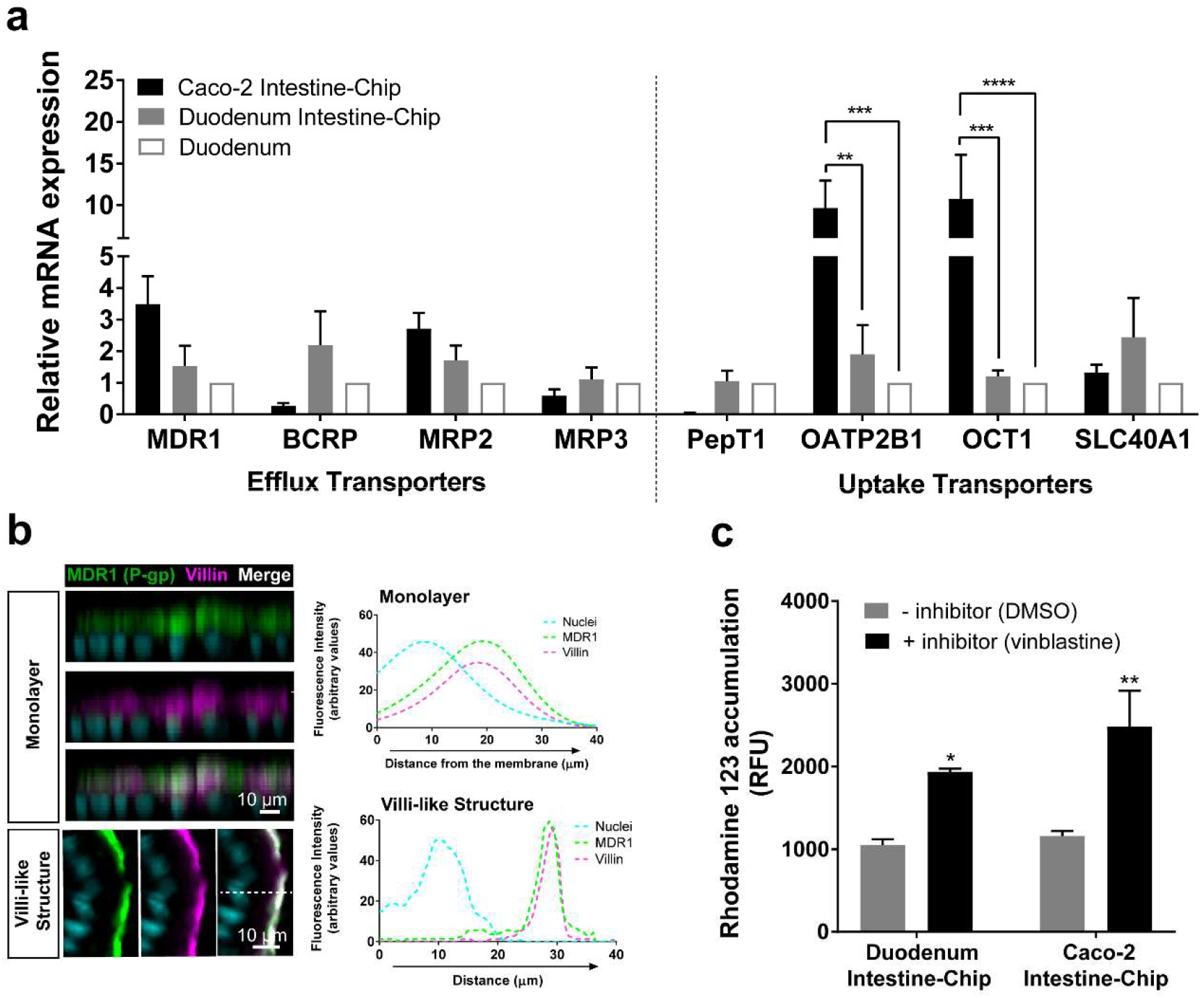
Duodenum Intestine-Chip shows the presence of major intestinal drug transporters and correct localization and function of efflux pump MDR1 (P-gp). **(a)** Comparison of the relative average gene expression levels of drug efflux (MDR1, BCRP, MRP2, MRP3) and uptake (PEPT1, OATP2B1, OCT1, SLC40A1) transporters in Caco-2 based Intestine-Chips (Intestine-Chip), three donor-specific Duodenum Intestine-Chips and RNA isolated directly from the duodenal tissue of three independent individuals (Duodenum). The results show that Duodenum Intestine-Chips express drug transport proteins at the levels close to human duodenal tissue. Note, that the expression of OATP2B1 and OCT1 in Caco-2 was significantly higher than in human duodenum while the difference between Duodenum Intestine-Chip and adult duodenum was not significant. Each value represents average gene expression ± s.e.m (error bars) from three independent experiments, each involving Duodenum Intestine-Chips established from a tissue of three independent donors (three chips/donor), RNA tissue from three independent biological specimens, and Caco-2 Intestine-Chips (three chips). Values are shown relative to the duodenal tissue expressed as 1, Two-way ANOVA, ****p<0.0001, ***p<0.001, **p<0.01. EPCAM expression was used as normalizing control. **(b)** Representative confocal immunofluorescence micrographs of vertical cross-sections (left) and corresponding to the plots (right) showing co-distribution of fluorescent signal of efflux transporter MDR1 (green), and villin (magenta) as visualized by merge channel (white) and fluorescent spectra overlap across vertical cross-sections of the differentiated epithelium in Duodenum Intestine-Chip. Efflux transporter MDR1 colocalize precisely with luminal cell surface marker (villin) in the monolayer of cells closely attached to membrane (monolayer) as well as in the cells lining subsequently formed villi-like structures (Villi-like Structure). Fluorescent signal representing cell nuclei is visualized in cyan. Scale bar, 10 µm. See also Figure 4—figure supplement 1 showing luminal localization of additional efflux (BCRP) and uptake (PEPT1) transporters in Duodenum Intestine-Chip. (c) Activity of efflux pump proteins. The intracellular accumulation of the fluorescent substrate of MDR1 - Rhodamine 123 was significantly increased in response to the MDR1 inhibitor vinblastine (black bars) in comparison to vehicle (DMSO) control (grey bars) in both systems – Caco2 and organoid-derived Duodenum Intestine-Chips. Data are presented as mean ± s.e.m (error bars) of at least three independent experiments involving chips generated from organoids of at least three individual donors and Caco-2 cell line. Two-way ANOVA, **p<0.01, *p<0.05.

We further demonstrated *in vivo* relevant localization of the luminal efflux pumps, MDR1, more commonly referred to as P-gp or P-glycoprotein (Figure 4B) and BCRP (Supplementary Figure 3A), as well as the uptake Peptide Transporter 1 (PEPT1) in organoids-derived Duodenum Intestine-Chips (Supplementary Figure 3B). All three transporters showed to co-distribute together with villin, a marker specific for apical cell membrane, at the intestinal cell brush border in Duodenum Intestine-Chip. This was confirmed on the cross-sectional confocal images of the duodenal epithelium cultured on chip as a co-localization (merge channel; white) between the fluorescent signal for MDR1, BCRP or PEPT1 and anti-villin staining. Similarly, plots depicting the distribution of the fluorescent signal across the cell Z-axis in these cross-sections revealed the presence of the significant overlap between the fluorescent spectra of each transporter and villin (Figure 4B and Supplementary Figure 3). Notably, the physiologically-relevant localization of MDR1, BCRP1 and PEPT1 at the luminal surface of intestinal epithelium was confirmed at two different time-points of Duodenum Intestine-Chip culture – when the cells have formed a confluent monolayer (around day 4 of culture) and after the villi-like structures morphogenesis has occurred (at day 8 of culture). The MDR1 activity was confirmed by measuring the intracellular accumulation of rhodamine 123 in the presence and absence of specific MDR1 inhibitor, vinblastine, across Duodenum Intestine-Chips and was compared to the expression in the Caco-2 Intestine-Chip model (Figure 4C). Addition of the inhibitor induced an ~2-fold increase in intracellular accumulation of rhodamine (1.84-fold increase in Duodenum Intestine-Chip and 2.14-fold increase in Caco-2 cells-based model), confirming active MDR1 efflux pumps in both cell systems.

### Drug-mediated CYP3A4 induction in the Duodenum Intestine-Chip

Induction of CYP450 drug metabolizing enzymes in human intestine is a major concern the pharmaceutical industry, as it is known to impact the pharmacokinetics and bioavailability of various orally-administered drugs, as well as, mediate drug-drug interactions. We therefore, evaluated the capability of our Duodenum Intestine-Chip to be applied for CYP3A4 induction studies and to help identify risk for drug-drug interactions in the clinic. This is not feasible in pre-clinical species such as rat and dog due to marked species differences in the expression and regulation of cytochromes P450 as well as substrate specificity of the nuclear receptors, such as pregnane X receptor (PXR), responsible for the transcriptional regulation of CYP3A4 and several drug transporters. Caco-2 Intestine-Chip has been previously reported to possess an increased activity of the CYP450 enzymes when compared to the conventional static culture of Caco-2 cells on transwell insert (25). However, the gene expression level of CYP3A4 measured in this system is significantly lower than in the adult human intestine (Figure 5A), limiting its application for pharmaceutical research, specifically pharmacokinetic evaluation. In the present study we demonstrate that, in comparison with the Caco-2 cells based system, our Duodenum Intestine-Chip expressed CYP3A4 at the much higher gene (~6000-times higher, p <0.0001) (Figure 5A) and protein level (Figure 5B), reaching the expression similar to the one observed in the adult human duodenum.

**Figure 5.**
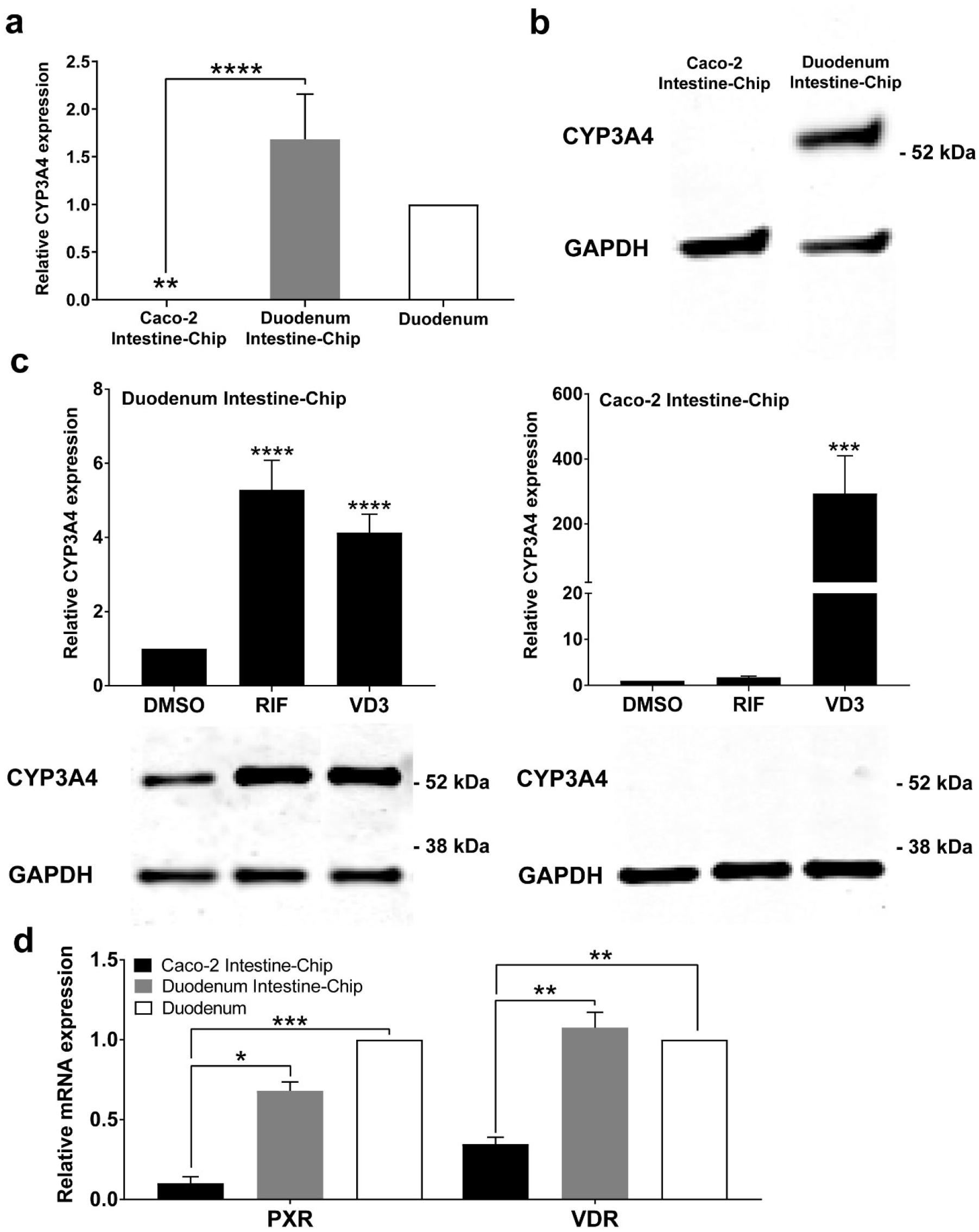
CYP3A4 expression levels and induction in Duodenum Intestine-Chip and Caco-2 cells-based Intestine-Chip. **(a)** Average gene expression levels of CYP3A4 ± s.e.m (error bars) in Caco-2 Intestine-Chip, Duodenum Intestine-Chip and human duodenum (three independent biological specimens). All values are shown relative to the adult duodenal tissue expressed as 1, One-way ANOVA, ****p<0.0001, **p<0.01. EPCAM expression was used as normalizing control. **(b)** Protein analysis of CYP3A4 in Caco-2 and Duodenum Intestine-Chips using western blot. **(c)** The CYP3A4 induction in Caco-2 Intestine-Chips and Duodenum Intestine-Chips treated with solvent (DMSO), 20 μM rifampicin (RIF) or 100 nM 1,25-dihidroxyvitamin D3 (VD3) for 48 h. The gene expression levels (top) of CYP3A4 were examined by real-time PCR analysis. On the y axis, the gene expression levels in the DMSO-treated chips were taken as 1.0. All data are represented as means ± s.e.m. Two-way ANOVA, ****p<0.0001 (compared with DMSO-treated cells). The corresponding CYP3A4 protein expression levels (bottom) were measured by western blotting analysis. **(d)** Gene expression analysis of the receptors PXR and VDR in organoids and Caco-2 cell-derived chips examined by real-time RT-PCR analysis and compared to their expression in adult duodenal tissue (Duodenum). On the y axis, the gene expression levels in adult tissue were taken as 1.0. All data are represented as means ± s.e.m. Two-way ANOVA, ***p<0.001, **p<0.01, *p<0.05.

In order to assess the drug-mediated CYP3A4 induction potential in the Duodenum Intestine-Chip, we exposed it to rifampicin (RIF) and 1,25-dihidroxyvitamin D3 (VD3), both of which are known prototypical CYP3A4 inducers. Treatment of Duodenum Intestine-Chips from all three independent donors with RIF and VD3 resulted in 5.3-fold and 4.1-fold induction of CYP3A4 expression, respectively, relative to that in the DMSO-treated controls (Figure 5C, left). Alternatively, Caco-2 Intestine-Chips were shown to respond to vitamin D3, but not RIF treatment (Figure 5C, right), which is consistent with the previously published reports ^47–49^. Dramatically low baseline expression of CYP3A4 in Caco-2 cells derived system showed to increase by 316-fold in the presence of VD3. However, it did not reach the levels observed in the human tissue and remained undetectable at the protein level as assessed by Western Blot analysis (Figure 5D, right). The differences between drug-mediated CYP3A4 induction potency of these two systems could be attributed to differences in gene expression levels of intestinal nuclear receptors, including PXR and vitamin D receptor (VDR). The gene expression levels of PXR and VDR in Duodenum Intestine-Chip were significantly higher and closer to human duodenal tissue than those in Caco-2 Intestine-Chip. An evident lack of PXR, which is known to be one of the key transcriptional regulators of CYP3A4 induction in humans, in Caco-2 cells may explain the differences observed between the Duodenum Intestine-Chip, and other Caco2–based systems including chips with Caco2. All together our findings demonstrate that the Duodenum Intestine-Chip represents a superior and more appropriate model for predicting CYP3A4-mediated drug-drug interactions in the intestine for oral drugs than Caco-2 cell-based models.

## Discussion

A number of *in vitro* and animal models have been developed and routinely applied for characterization of distribution and prediction of absorption, metabolism, and excretion (ADME) of xenobiotics in humans. Among these, one of the most widely used models is the conventional Caco-2 monolayer culture, considered as the current “gold standard” in studying intestinal disposition of drugs *in vitro*. However, inherent limitations, such as lack of *in vivo* relevant 3D cytoarchitecture, lack of appropriate ration of cell populations, altered expression profiles of drug transporters and drug metabolizing enzymes, especially CYP450s, and aberrant CYP450 induction response, challenge the use of these model for predicting ADME in the clinic. On the other side of the spectrum, animal models, while retaining proper physiological conditions, exhibit species differences in both drug metabolism and drug transport, as well as substrate specificity for nuclear receptors regulating CYP450s and transporters. For these reasons, there is a need for new systems for predicting human ADME and determining risk for drug-drug interactions mediated by intestinal CYP450s and drug transporters.

Current preclinical models are not able to fully recapitulate the complex nature and function of human intestinal tissue, leading to a limited accuracy and poor predictability in drug development. In this study, we leveraged Organs-on-Chips technology and primary organoids to emulate multicellular complexity, physiological environment, native intestinal tissue architecture and functions and to create an alternative human-relevant model for the assessment of ADME of orally administered drugs. Similarly, to our previous study ^31^ morphological analysis of the Duodenum Intestine-Chip confirmed the establishment of intact tissue-tissue interface formed by organoids-derived epithelium and small intestinal microvascular endothelial cells. The application of the mechanical stimulation in the form of continuous luminal and vascular flow showed to exert the beneficial effect on the intestinal tissue architecture. Increased epithelial cell height, acquisition of cobblestone-like cell morphology, formation of well-defined cell-cell junctions, and dense intestinal microvilli was attributed to the presence of flow. We confirmed the development of comparable levels of intestinal barrier function to hydrophilic solute (dextran) across Duodenum Intestine-Chip cultures established from organoids, which were isolated from three independent donors. This functional feature of intestinal tissue becomes critical when studying drug uptake, efflux, and disposition in polarized organ systems. Although many studies have demonstrated that Caco-2 transwell allow good prediction of transcellular drug absorption, they showed to be unreliable in the assessment of passive diffusion of polar molecules such as hydrophilic drugs and peptides. This is related to much smaller, in respect to small intestinal villi, effective area of the monolayer and its higher tight junctional resistance ^50, 51^. Organoids have quickly become a model of interest in drug discovery and development applications, yet the use of organoids for the studies of intestinal permeability has been linked with major technical difficulties related to their three-dimensional structure, limited access to lumen, and the need for the use of sophisticated microinjection techniques for the application of the compound at the apical cell surface. Therefore, Duodenum Intestine-Chip that emulates complex intestinal tissue architecture, as evidenced by the presence of three-dimensional villi-like structures on scanning electron micrographs acquired on day 8 of fluidic culture, and allows a direct exposure of apical cell surface to test compounds represents a major technological advantage over Caco-2 transwell and organoids and might provide a valid alternative for the preclinical studies of intestinal drug absorption.

Immunofluorescent imaging and gene expression analysis demonstrated that Duodenum Intestine-Chip possess specialized intestinal cell subpopulations, that are absent in tumor-derived cell lines including Caco-2. Importantly, they are present in the chip at the physiologically relevant ratios, similar to the ones observed in the native human duodenum. Expression of the markers specific for the intestinal types that naturally reside in the villus compartment – alkaline phosphatase for absorptive enterocytes, mucin 2 for goblet cells, chromogranin A for enteroendocrine cells – showed to increase up to day 8 of Duodenum Intestine-Chip culture. This has been accompanied by the decline in the expression of genes specific for crypt-residing cells – lysozyme for Paneth cells and Lgr5 for stem cells (data not shown) suggesting the acquisition of terminally differentiated cell phenotype by all of the cells present in the chip. This is in line with well-known effect of Wnt3a withdrawal on organoids-derived primary intestinal epithelium ^20, 21^. Because human intestinal cells do not produce significant amounts of Wnt3A, EGF, or several other growth factors essential for stem cell division and cell proliferation, removal of Wnt3A supplementation leads to a loss of LGR5-positive cells, decreased cell proliferation, appearance of secretory cell lineages, including goblet and enteroendocrine cells, and the transformation of immature crypt-like enterocytes into differentiated nutrient-absorptive cells. The presence of differentiated cell subpopulations is critical for modelling various aspects of intestinal biology and function, including mucus production, antimicrobial response, host-microbiome interaction, secretion of intestinal hormones, nutrients digestion and absorption. Given that we have shown that the Duodenum Intestine-Chip recapitulates faithfully the physiologically relevant ratios of these cells it could be readily applied beyond ADME applications in development of new approaches and therapeutics that targets specific cell populations. For example, it could serve to test to target relevant biology of Paneth cells, which through a series of genetic studies have been implicated in inflammatory bowel disease ^52–55^.

We compared global RNA expression profiles, obtained by RNA-seq, of Duodenum Intestine-Chip, duodenal organoids and human native duodenal tissue in order to determine which of the two models better reflects their natural counterpart. Surprisingly, although chips and organoids were established with the cells of the same origin (donors, source of the tissue) their transcriptomic profiles were shown to significantly differ from each other – the total number of 472 up- and downregulated genes were found in this comparison. Moreover, several different analyses demonstrated that the transcriptome of Duodenum Intestine-Chip more closely resembles global gene expression in human adult duodenum than do the organoids, strongly suggesting that Duodenum Intestine-Chip constitute a more accurate representation of human *in vivo* tissue. Importantly, the subset of 305 genes that were found to be common for Duodenum Intestine-Chip and human tissue but different from organoids, showed to be associated with important biological functions including: digestion, transport of nutrients and ions, extracellular matrix organization, wound healing, metabolism, detoxification as well as tissue development. In addition, several genes involved in drug metabolism, including but not limited to drug metabolism enzymes: CYP3A4, UGT1A4, UGT2A3, UGT2B7, UGT2B15, UGT2B17, were observed to exhibit similar pattern of expression in Duodenum Intestine-Chip and adult duodenal tissue while they differed from organoids, suggesting the improved potential of this system to study biotransformation of xenobiotics and drug-drug interactions.

Intestinal efflux and uptake transporters are key determinants of absorption and subsequent bioavailability of a large number of orally administered drugs. *In vivo*-like expression of the major intestinal drug transporters, including clinically relevant MDR1, BCRP and PEPT1, was demonstrated in Duodenum Intestine-Chip system. Additionally, we confirmed the apical localization, which is known to be crucial for the unique gatekeeper function of these proteins in controlling drug access to metabolizing enzymes and excretory pathways ^1, 56^, within the plasma membrane of intestinal epithelial cells grown on chip. Functional assessment of MDR1 efflux revealed similar level of activity as observed in Caco-2 Intestine-Chip and its successful inhibition by vinblastine. Notably, in comparison to Caco-2 Intestine-Chip the organoids-based chip system showed improved relative mRNA expression levels of organic anion and cation transporters: OATP2B1 and OCT1. These transporters are responsible for the uptake of numerous xenobiotics, including statins, antivirals, antibiotics, and anticancer drugs ^57^. While significantly higher expression of these proteins, in comparison to human duodenal tissue, was observed in Caco-2 model, their levels showed to be closer to *in vivo* in Duodenum Intestine-Chip. Our findings suggest that organoids-derived Intestine-Chip system could be applied to assess the specific contribution of efflux transporters to drug disposition, to evaluate the active transport of xenobiotics across intestinal barrier, as well as, for the modelling of increased absorption by targeting of specific uptake transporters such as PEPT1 or OATP2B1.

Evaluation of the expression and drug-mediated induction of the CYP3A4, a major enzyme involved in human metabolism of xenobiotics revealed a key advantage of Duodenum Intestine-Chip over Caco-2 models. Significantly higher and much closer to human duodenal tissue expression of CYP450 enzyme was observed in Duodenum Intestine-Chip in comparison to Caco-2 cells-based system. Consistent with the results of others ^47^ CYP3A4 was undetectable at the protein level in Caco-2 cells and remained unchanged upon stimulation with RIF (a PXR agonist). While successful and reproducible modulation of CYP3A4 expression using agonists specific for PXR and VDR, rifampicin and vitamin D3, respectively, was achieved in Duodenum Intestine-Chips engineered individually from the organoids of three independent donors. CYP3A4 induction levels observed were similar to those shown for intestinal slices ^58^, suggesting suitability of this model for the studies of drug metabolism, that cannot be supported by the previous Caco2 models.

In conclusion, Duodenum Intestine-Chip provides a faithful representation of human duodenum and represents a potential tool for preclinical drug assessment in human-relevant system. Moreover, as it’s composed of the cells isolated from individual patient it could in the future be personalized in order to assess interindividual differences in drug disposition and responses, studies of the effect of genetic polymorphisms on pharmacokinetics and pharmacodynamics as well as decoupling of the effect of various factors such as age, sex, disease state and diet on metabolism, clearance and bioavailability of xenobiotics. This system could also be applied help us to better understand the basic biology of human intestinal tissue in health and disease state and potentially enable novel therapeutic development as we further our understanding of mechanisms driving key disease phenotypes.

## Materials and Methods

### Human tissue collection, generation, and culture of organoids

Human duodenal organoids cultures were established from biopsies obtained during endoscopic or surgical procedures utilizing the methods developed by the laboratory of Dr. Hans Clevers^21^. De-identified biopsy tissue was obtained from healthy adult subjects who provided informed consent at Johns Hopkins University and all methods were carried out in accordance with approved guidelines and regulations. All experimental protocols were approved by the Johns Hopkins University Institutional Review Board (IRB #NA 00038329). Briefly, organoids generated from isolated intestinal crypts were grown embedded in Matrigel™ (Corning, USA) in the presence of Expansion Medium consisting of Advanced DMEM F12 supplemented with 50% v/v Wnt3A conditioned medium (produced by L-Wnt3A cell line, ATCC CRL-2647), 20% v/v R-spondin-1 conditioned medium (produced by HEK293T cell line stably expressing mouse R-spondin1; kindly provided by Dr Calvin Kuo, Stanford University, Stanford, CA), 10% v/v Noggin conditioned medium (produced by HEK293T cell line stably expressing mouse Noggin), 10mM HEPES, 0.2 mM GlutaMAX, 1x B27 supplement, 1x N2 supplement, 1 mM n-acetyl cysteine, 50 ng/ml human epithelial growth factor, 10 nM human [Leu15]-gastrin, 500 nM A83-01, 10 μM SB202190, 100 μg/ml primocin. EM was replaced every other day and supplemented with 10 μM CHIR99201 and 10 μM Y-27632 during the first 2 days after passaging. Organoids were passaged every 7 days and used for chip seeding between passage numbers 5 and 30.

### Duodenum Intestine-Chip

The design and fabrication of Organ-Chips used to develop the Duodenum Intestine-Chip was based on previously described protocols^59^. The chip is made of a transparent, flexible polydimethylsiloxane (PDMS), an elastomeric polymer. The chip contains two parallel microchannels (a 1 × 1 mm epithelial channel and a 1 × 0.2 mm vascular channel) that are separated by a thin (50 μm), porous membrane (7 μm diameter pores with 40 μm spacing) coated with ECM (200 µg/ml collagen IV and 100 µg / ml Matrigel™ at the epithelial side and 200 µg / ml collagen IV and 30 µg / ml fibronectin at the vascular side). Chips were seeded with intestinal epithelial cells obtained from enzymatic dissociation of organoids, as described previously^31^, and incubated overnight before being washed with fresh media. Next day, the chips were connected to the culture module instrument (inside the incubator), that can hold up to 12 chips and allows for control of flow and stretching within the chips using pressure driven flow^60^. The chips are maintained under constant perfusion of fresh expansion medium at 30 µl / hr through top and bottom channels of all chips until day 6. Human Intestinal Microvascular Endothelial Cells (HIMECs; Cell Biologics) were than plated on the vascular side of the ECM-coated porous membrane in EGM2-MV medium, which contains human epidermal growth factor, hydrocortisone, vascular endothelial growth factor, human fibroblastic growth factor-B, R3-Insulin-like Growth Factor-1, Ascorbic Acid and 5% fetal bovine serum (Lonza Cat. no. CC-3202). At this time the medium supplying the epithelial channel was switched to differentiation medium. Differentiation medium consisted of the same components as those of Expansion Medium with 50% less Noggin and R-spondin and devoid of Wnt3A and SB202190. Media supplying both chip channels were under continuous flow. Cyclic, peristalsis-like deformations of tissue attached to the membrane (10% strain; 0.2 Hz) were initiated after the formation of confluent monolayer at ~4 days in culture.

### Permeability assays

In order to evaluate the establishment and integrity of the intestinal barrier, 3kDa Cascade Blue dextran (Sigma, D7132) was added to the epithelial compartment of the Duodenum Intestine-Chip at 0.1 mg / ml on the day of their connection to flow. Effluents of the endothelial compartment were sampled every 48 hours to determine the concentration of dye that had diffused through the membrane. The apparent paracellular permeability (Papp) was calculated based on a standard curve and using the following formula:

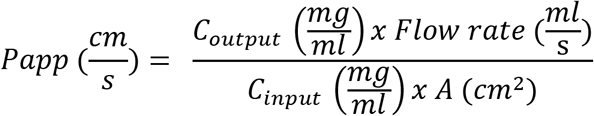

Where *C*_*output*_ is the concentration of dextran in the effluents of the endothelial compartment, *A* is the seeded area, and *C*_*input*_ is the input concentration of dextran spiked into the epithelial compartment. The establishment of intestinal barrier function in Duodenum Intestine-Chips was evaluated in three independent experiments, each performed using chips established from a different donor of biopsy-derived organoids. At least three different chips were used per condition.

### Morphological analysis

Immunofluorescent staining of cells in the Organ-Chip was performed with minor modifications to previously reported protocols^32^. Cells were fixed with 4% formaldehyde or cold methanol, and, when required, were permeabilized, using 0.1% Triton X-100. 5% (v/v). Donkey serum solution in PBS was used for blocking. Incubation with primary antibodies directed against ZO-1, VE-cadherin, E-cadherin, villin, mucin 2, lysozyme, chromogranin A, MDR1, BCRP, PEPT1 (see Supplementary Table 2) was performed overnight at 4°C. Chips treated with corresponding Alexa Fluor secondary antibodies (Abcam) were incubated in the dark for 2 hours at room temperature. Cells were then counterstained with nuclear dye DAPI. Images were acquired with an inverted laser-scanning confocal microscope (Zeiss LSM 880 with Airyscan).

Chips processed for SEM were fixed in 2.5% glutaraldehyde, treated with 1% osmium tetroxide in 0.1 M sodium cacodylate buffer, dehydrated in a series of graded concentrations of ethanol solutions and critical point dried, as described previously^31^. Prior to imaging, samples were coated with a thin (10 nm) layer of Pt/Pd using a sputter coater.

### Measurement of the density of microvilli

Images of microvilli on the surface of cells were captured with a scanning electron microscope (JSM-5600LV; JEOL). The morphological analysis and quantification of microvilli was performed using ImageJ. Number of intestinal microvilli per µm^2^ were calculated after applying two image processing techniques, namely binarization and particle analysis, and Otsu’s thresholding method, as described previously ^61^.

### MDR-1 efflux pump activity

The transporter activity of MDR-1 was assessed using the MDR1 Efflux Assay Kit (ECM910, Millipore), as per manufacturer’s instructions. Briefly, Duodenum Intestine-Chips and Caco-2 Intestine-Chips^33, 62^ were perfused apically with rhodamine 123, a fluorescent transport substrate of MDR1. Intracellular accumulation of dye, was detected by fluorescent imaging (Olympus IX83) and measured in the presence and absence of MDR-1 specific inhibitor vinblastine (22 µM). Three independent experiments were performed for Caco-2 Intestine-Chips and Duodenum Intestine-Chips, each using chips established from a different donor of biopsy-derived organoids. At least three different chips were used per condition. Images of each were taken at three different fields of view, digitally processed and quantified using Fuji software.

### CYP3A4 induction

Duodenum Intestine-Chips and Caco-2 Intestine-Chips were treated with 100 nM 1,25-Dihydroxyvitamin D3 (Sigma) or 20 µM rifampicin (Sigma), which are known to induce CYP3A4, for 48 hours. Controls were treated with DMSO (final concentration 0.1%). Subsequently, cells in the epithelial channel were harvested either for RNA isolation and gene expression analysis or for Western blotting in order to assess the level of CYP3A4 induction at the gene and protein level, respectively.

### Western blotting

RIPA cell lysis buffer (Pierce) supplemented with protease and phosphatase inhibitors (Sigma) was used for the extraction of total protein from the chips. The protein concentration in each sample was determined using the bicinchoninic acid method. Equal amounts (15 µg) of protein lysates were heat denatured and separated on a 4–10% Mini-Protean Precast Gel (Bio-Rad), followed by transfer on a nitrocellulose membrane (Bio-Rad). After blocking with 5% nonfat milk, membranes were probed with primary antibodies for CYP3A4 (mouse monoclonal, Santa Cruz Biotechnology) and GAPDH (rabbit polyclonal, Abcam) and incubated overnight at 4°C, followed by incubation for 1 hour with IRDye-conjugated secondary antibodies against rabbit and mouse immunoglobulin G (LI-COR), at room temperature. Finally, blots were scanned using an Odyssey Infrared Imaging System (LI-COR) and the protein bands were visualized and quantified using Image Studio software (LI-COR). Gluceraldehyde-3-phosphatase dehydrogenase (GAPDH) was used as the loading control.

### Gene expression analysis

Total RNA was isolated from the chip using PureLink RNA Mini kit (Fisher Scientific) and reverse transcribed to cDNA using SuperScript IV Synthesis System (Fisher Scientific). qRT-PCR was performed using TaqMan Fast Advanced Master Mix (Applied Biosystems) and TaqMan Gene Expression Assays (see Supplementary Table 3, Fisher Scientific) in QuantStudio 5 PCR System (Fisher Scientific). Relative expression of gene was calculated using 2-∆∆Ct method.

### RNA isolation and sequencing

RNA was extracted using TRIzol (Life Technologies) according to manufacturer’s guidelines. Samples were submitted to GENEWIZ South Plainfield, NJ for next generation sequencing. After quality control and a complementary DNA library creation, all samples were sequenced using HiSeq 4000 with 2×150 bp paired-end reads per sample.

### RNA sequencing bioinformatics

Pre-processing: raw sequence data (.bcl files) generated from Illumina HiSeq was converted into fastq files and de-multiplexed using Illumina’s bcl2fastq 2.17 software. Read quality was assessed using FastQC. Adaptor and low-quality (< 15) sequences were removed using Trimmomatic v.0.36. The trimmed reads were mapped to the Homo sapiens reference genome available on ENSEMBL using the STAR aligner v.2.5.2b. The STAR aligner uses a splice aligner that detects splice junctions and incorporates them to help the alignment of the entire read sequences. BAM files were generated following this step. Unique gene raw counts were calculated by using feature Counts from the Subread package v.1.5.2. Only the unique reads that fell within exon regions were counted.

Differential Gene Expression Analysis: to combine our gene expression dataset with the publicly available data for adult duodenum gene expression (provided as Fragments Per Kilobase of transcript per Million mapped read (FPKM) values in ^63^), we converted the raw counts to FPKMs. Then using the log2(FPKM) expressions of the combined datasets, we applied DE gene analysis using the R package “limma”^35^. For each comparison, the thresholds used to identify the DE genes were set to a) adjusted p-values < 0.05 and b) absolute log2 fold change > 2.

### GO term enrichment analysis and KEGG Pathway analysis

After expression pattern clustering, the transcripts from specific groups were subjected to functional annotation, including GO (Gene Ontology) functional annotation and KEGG (Kyoto Encyclopedia of Genes and Genomes) pathway annotation. The GO terms and KEGG pathway enrichment was performed using The Database for Annotation, Visualization and Integrated Discovery (DAVID v 6.8, http://david.abcc.ncifcrf.gov).

### Statistical analysis

All experiments were performed in triplicates and repeated with organoids from three different human donors. One-way or two-way ANOVA was performed to determine statistical significance, as indicated in the figure legends. The error bars represent standard error of the mean [s.e.m]; P values < 0.05 and above were considered as significant.

### Data availability

RNA sequencing data have been deposited in the National Center for Biotechnology Information Gene Expression Omnibus (GEO) under accession number GSE135196.

## Supporting information

Figure3- source data 1. Differentially Expressed Genes in Duodenum Intestine-Chip vs. Duodenal Organoids, Related to Figure 3.

Figure3- source data 2. Differentially Expressed Genes in Adult Duodenum vs. Duodenal Organoids, Related to Figure 3.

Figure3- source data 3. Differentially Expressed Genes Common in Duodenum Intestine-Chip and Adult Duodenum versus Duodenal Organoids, Related to Figu

Figure3- source data 4. Enriched GO Terms from a List of Differentially Expressed Genes Common in Duodenum Intestine-Chip and Adult Intestine versus D

Supplementary Figure 2 - source data 1. Differentially Expressed Genes in Duodenal Organoids vs. Adult Duodenum, Related to Supplementary Figure 2

Supplementary Figure 2 - source data 2. Differentially Expressed Genes in Duodenum Intestine-Chip vs. Adult Duodenum, Related to Supplementary Figure 2

---

**Figure 4—figure supplement 1.**
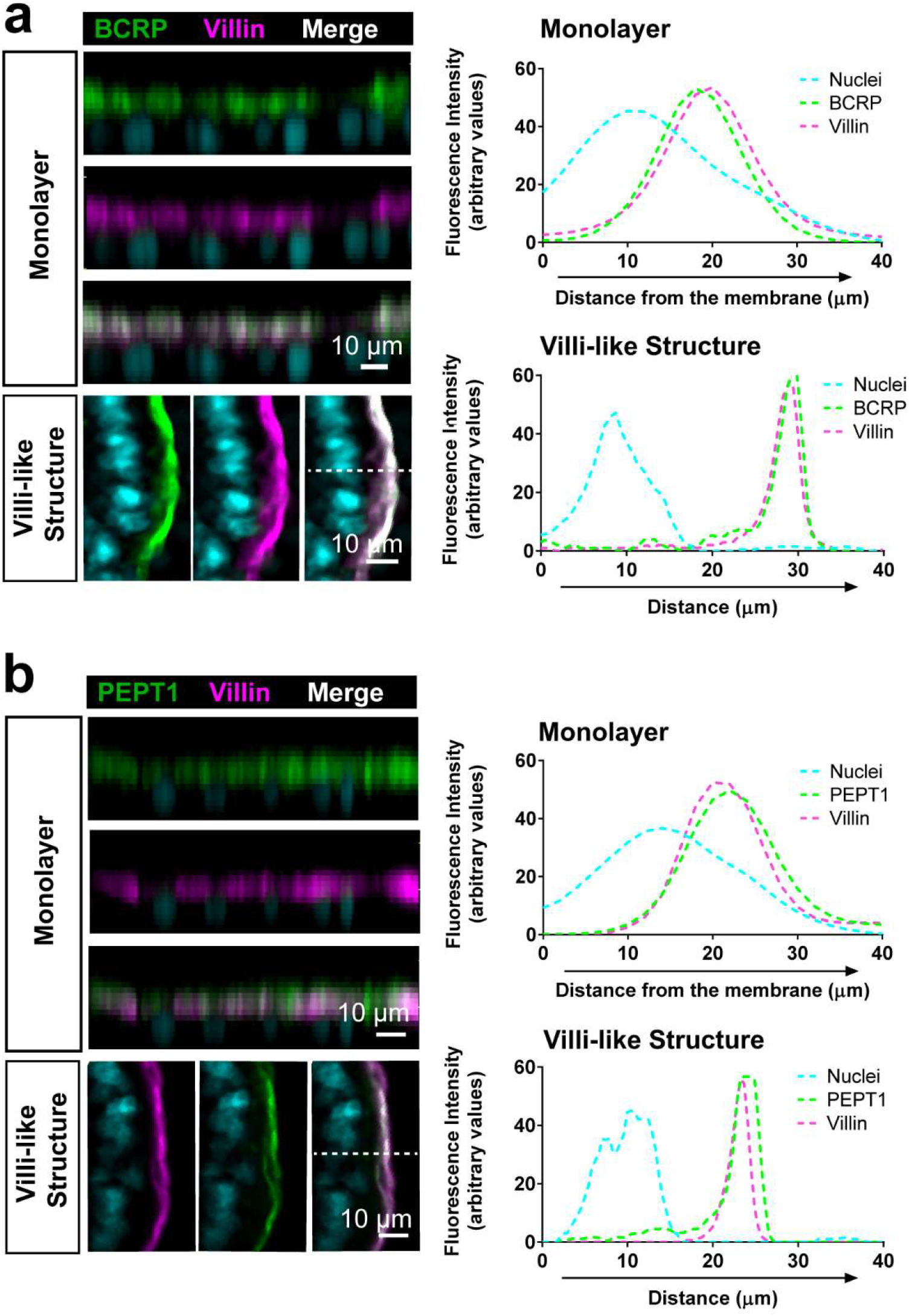
Luminal localization of efflux (BCRP) and uptake (PEPT1) transporters in Duodenum Intestine-Chip. Representative cross-sectional confocal images of Duodenum Intestine-Chip (left) showing apical localization of efflux BCRP (**a**; green) and uptake PEPT1 drug transporters (**b**; green) that co-localize with luminal cell surface marker villin (magenta) at the time of confluent monolayer formation as well as within successively formed villi-like structures. Plots representing distribution of the fluorescence signal across epithelial cell z-axis revealed close overlap of green (transporters; BRCP and PEPT1) and magenta (apical cell marker; villin) signals confirming co-distribution of these proteins on the luminal cell surface. Cell nuclei are visualized in cyan. Scale bar, 10 µm.

**Supplementary Figure 1.**
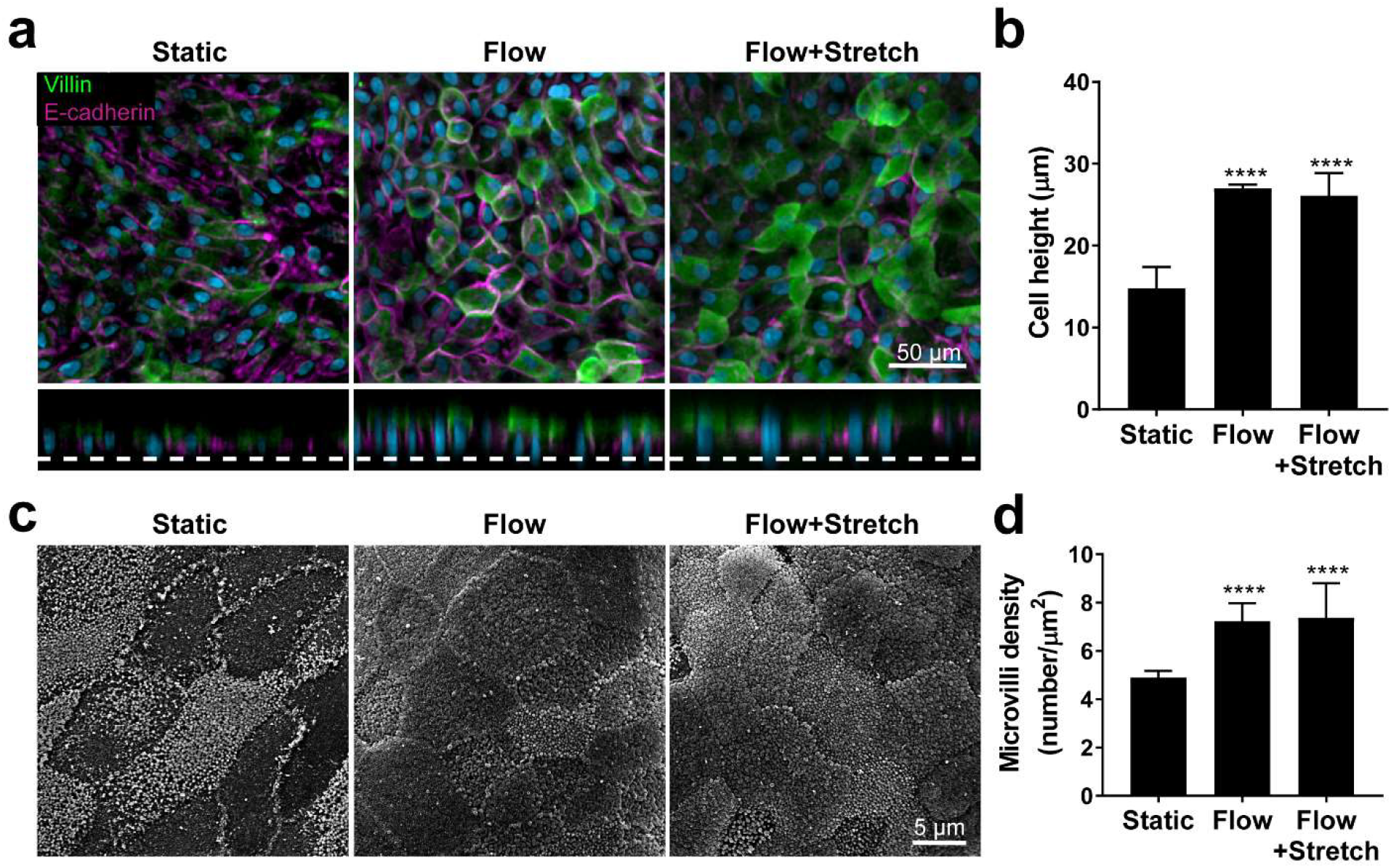
Flow-induced increase in primary intestinal epithelial cells height and microvilli formation. **(a)** Representative confocal images of x-y (top) and x-z (bottom) optical sections of duodenal organoid-derived epithelial cells cultured under static (Static) or fluid flow (30 µl h^−1^; Flow) or flow and stretch (30 µl h^−1^; 10% strain, 0.2 Hz; Flow+Stretch) conditions and stained for apical marker villin (green) and basolateral protein E-cadherin (magenta). Nuclei we counterstained with DAPI (grey). Scale bar, 50 µm. **(b)** The quantitative analysis of the average cell height measured from Z-stack images as the distance between apical marker villin (green, a) and PDMS membrane (dotted line, a). The data represent the mean ± s.e.m; One-way ANOVA, ****p<0.0001. **(c)** Scanning electron microscopy surface images of duodenal enteroid-derived epithelium cultured under static or flow +/− stretch conditions. Cells seeded in the top channel of Organ-Chip were maintained with (Flow) or without (Static) medium perfusion (30µl h^−1^) in both channels and 10% of mechanical stretch (0.2 Hz) (Flow+Stretch). Images were captured at the center area of the chamber. Scale bar, 5 µm. **(d)** Quantification of microvilli. Density of microvilli per µm^2^ was measured from the SEM images (100 µm2, 20 FOV) as described in the Methods. The data represent the mean ± s.e.m; One-way ANOVA, ****p<0.0001.

**Supplementary Figure 2.**
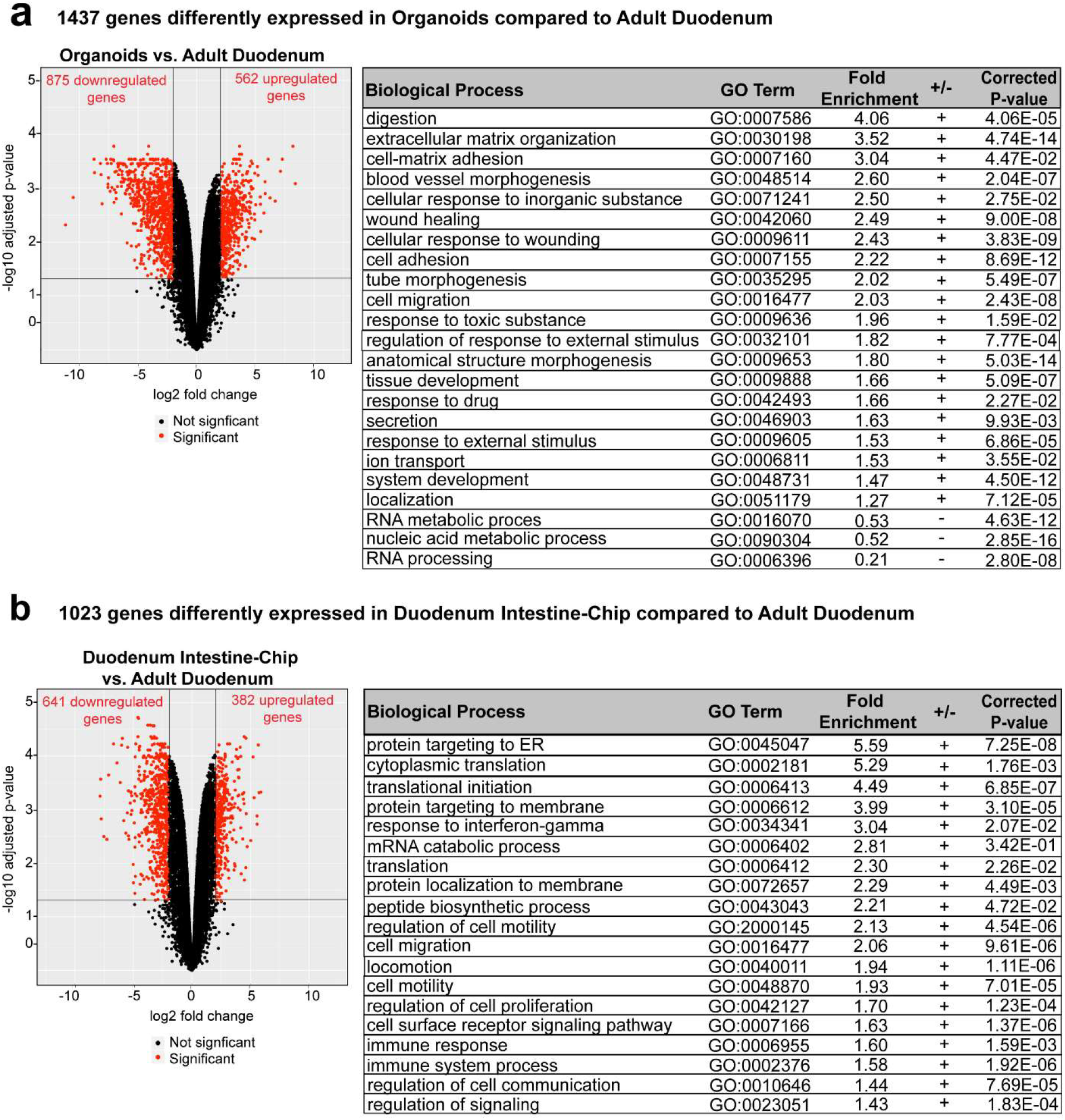
Differentially expressed genes and enriched Gene Ontology categories in Organoids or Duodenum Intestine-Chip with respect to Adult Duodenum. **(a)** Volcano plot (left) and functional enrichment analysis (right) of differentially expressed genes between Organoids and Adult Duodenum (Supplementary Figure 2—source data 1). The red dots represent genes that are significantly (adj. p-value<0.05) up- or downregulated. The black dots correspond to the non-differentially expressed genes. The vertical lines correspond to 2.0-fold up and down and the horizontal line has been drawn at the level of the selected cutoff adjusted p-value (adj. p-value<0.05). Sample sizes were as follows: Duodenum Intestine-Chip, n =3; Organoids, n = 3. All samples were biologically independent (derived from a different donor). Samples from the same 3 donors were used for the establishment of Duodenum Intestine-Chip and organoid cultures. Functional enrichment analysis demonstrated over (+) and under (-) represented biological processes in the GO categories concerning digestion, extracellular matrix organization, angiogenesis, cell adhesion, tissue development, cell response to drugs and toxic substances, while nucleic acid metabolic process and RNA processing were under represented. GO, Gene Ontology. **(b)** Differential gene expression and functional enrichment analysis between Duodenum Intestine-Chip and adult human tissue (Supplementary Figure 2—source data 2) demonstrating up- and down-regulated genes (volcano plot, left) and annotated to them biological processes (table, right) involving but not limited to protein synthesis and targeting as well as cell cycle and cell proliferation. Red dots: significant genes (adj. p-value<0.05). Black dots: non-differentially expressed genes. Sample sizes were as follows: Duodenum Intestine-Chip, n =3; Adult Duodenum Human tissue, n = 2. Sample sizes were as follows: Duodenum Intestine-Chip, n =3; Adult Duodenum, n = 2. All samples were biologically independent (derived from a different donor). **Supplementary Figure 2—source data 1.** Differentially Expressed Genes in Duodenal Organoids vs. Adult Duodenum, Related to Supplementary Figure 2. (added as Supplementary Materials) **Supplementary Figure 2—source data 2.** Differentially Expressed Genes in Duodenum Intestine-Chip vs. Adult Duodenum, Related to Supplementary Figure 2. (added as Supplementary Materials)

**Supplementary Table 1.** RNAseq Datasets Downloaded from Public Databases

| Sample Label | Description | Source | Donor ID | Accession # |
| --- | --- | --- | --- | --- |
| Adult Duo_1 | Adult Duodenum | EMBL-EBI<br>ArrayExpress | V145 | E-MTAB-1733 (duodenum_4b) |
| Adult Duo_2 | Adult Duodenum | EMBL-EBI<br>ArrayExpress | V150 | E-MTAB-1733 (duodenum_4c) |

**Supplementary Table 2.** List of primary antibodies used for immunostaining.

| Target Protein | Host | Vendor | Catalog Number | Dilution |
| --- | --- | --- | --- | --- |
| BCRP | Mouse | EMD Millipore | MAB4155 | 1:100 |
| Chromogranin A | Goat | Santa Cruz Biotechnology | sc-1488 | 1:100 |
| E-Cadherin | Mouse | Abcam | ab1416 | 1:100 |
| Lysozyme | Rabbit | Dako | A0099 | 1:500 |
| MDR-1 (P-gp) | Mouse | Thermo Fisher | MA5-13854 | 1:100 |
| Mucin-2 | Mouse | Santa Cruz Biotechnology | sc-7314 | 1:400 |
| PEPT1 | Mouse | Santa Cruz Biotechnology | sc-373742 | 1:100 |
| VE-Cadherin | Rabbit | Abcam | ab232880 | 1:400 |
| Villin | Rabbit | Abcam | ab130751 | 1:100 |
| ZO-1 | Mouse | Abcam | ab61357 | 1:100 |

**Supplementary Table 3.** List of human TaqMan Gene Expression Assays used for qRT-PCR.

| Gene | Gene Name | TaqMan Assay ID |
| --- | --- | --- |
| ALPI | Alkaline Phosphatase | Hs00357579_g1 |
| BCRP/ ABCG2 | ATP binding cassette subfamily G member 2 | Hs01053790_m1 |
| CHGA | Chromogranin A | Hs00900375_m1 |
| CYP3A4 | Cytochrome family 3 subfamily A member 4 | Hs00604506_m1 |
| EpCAM | Epithelial Cell Adhesion Molecule | Hs00901885_m1 |
| LYZ | Lysozyme | Hs00426232_m1 |
| MDR1/ABCB1 | ATP binding cassette subfamily B member 1 | Hs00184500_m1 |
| MRP2/ ABCC2 | ATP binding cassette subfamily C member 2 | Hs00960489_m1 |
| MRP3/ ABCC3 | ABCC3 | Hs00978452_m1 |
| MUC2 | Mucin 2 | Hs00894025_m1 |
| OAT2B1/ SLCO2B1 | Solute carrier organic anion transporter family member 2B1 | Hs01030343_m1 |
| OCT1/ SLC22A1 | Solute carrier family 22 member 1 | Hs00427552_m1 |
| PEPT1/SLC15A1 | Solute carrier family 15 member 1 | Hs00192639_m1 |
| PXR/ NR1I2 | Nuclear receptor subfamily 1 group I member 2 | Hs01114267_m1 |
| SLC40A1 | Solute carrier family 40 member 1 | Hs00205888_m1 |
| VDR | Vitamin D (1,25- dihydroxyvitamin D3) receptor | Hs01045843_m1 |

